# Interplay between BALL and CBP maintains H3K27 acetylation on active genes in *Drosophila*

**DOI:** 10.1101/2021.07.02.450821

**Authors:** Muhammad Haider Farooq Khan, Ammad Shaukat, Zain Umer, Hina Ahmad, Muhammad Tariq

## Abstract

CREB binding protein (CBP) is a multifunctional transcriptional co-activator that interacts with a variety of transcription factors and acts as a histone acetyltransferase. In *Drosophila*, CBP mediated acetylation of histone H3 lysine 27 (H3K27ac) is a known hallmark of gene activation regulated by trithorax group proteins (trxG). Recently, we have shown that a histone kinase Ballchen (BALL) substantially co-localizes with H3K27ac at trxG target loci and is required to maintain gene activation in *Drosophila*. Here, we report direct interaction between BALL and CBP, which positively regulates H3K27ac. Analysis of genome-wide binding profile of BALL and CBP reveals major overlap and their co-localization at actively transcribed genes. We show that BALL biochemically interacts with CBP and depletion of BALL results in drastic reduction in H3K27ac. Together, these results demonstrate a previously unknown synergy between BALL and CBP and reveals a potentially new pathway required to maintain gene activation during development.

## Introduction

In metazoans, covalent modifications of histones act in a combinatorial manner to regulate developmental gene expression (1, 2). Among these modifications, histone H3 lysine 27 acetylation (H3K27ac) is considered a hallmark of trithorax group (trxG) mediated gene activation that antagonizes repression by Polycomb group (PcG) proteins (3). Although the anti-silencing effect of H3K27ac and its catalyzing enzyme CBP is widely documented in flies and mammals, how this CBP mediated H3K27ac is maintained on actively transcribed genes remains elusive (3, 4). In *Drosophila*, presence of only one homolog of CBP known as *nejire*, makes it suitable to understand its function. Importantly, chromatin immunoprecipitation sequencing (ChIP-seq) data from previously published reports revealed the presence of CBP on a majority of actively transcribed genes in *Drosophila* S2 cells (5). Our lab recently showed that *Drosophila* Ballchen (BALL), a known serine-threonine kinase that mediates histone H2AT119 phosphorylation (6), associates with trxG target genes enriched with H3K27ac (7). In this report, we have further analyzed the link between BALL, H3K27ac and CBP. We have discovered that BALL not only binds to chromatin enriched with H3K27ac but also shares more than 77% genomic binding sites with CBP. Both BALL and CBP primarily associate with transcription start sites (TSS) and regions up to 1Kb upstream of TSS. Four of top five DNA motifs that CBP binds are also bound by BALL. Importantly, BALL biochemically interacts with CBP and positively contributes to maintenance of H3K27ac by CBP since depletion of BALL leads to diminished H3K27ac.

## Results and Discussion

Recently we reported that BALL exhibit trxG like behavior by contributing to the maintenance of gene activation in flies (7). Since BALL was shown to co-localize with Trithorax (TRX) at genes enriched with H3K27ac, we investigated if BALL binds to chromatin together with CBP. To this end, we analyzed ChIP-seq data of BALL (7) and CBP (5) from *Drosophila* S2 cells. Comparison of BALL and CBP binding to chromatin revealed their co-occupancy at 4577 genes (Fig. 1*A* and Dataset S1). Further analysis revealed that both BALL and CBP shared binding sites in genomic regions up to 1Kb upstream of the transcription start sites (TSS), however, CBP is also found at the regions that are further upstream of TSS (Fig. 1*B* and *C*). This finding is in line with the reported binding of CBP in promoter as well as enhancer regions (5, 8). Analysis of the genomic distribution of CBP and BALL in terms of their binding at promoter regions, within gene body, introns and intergenic regions also revealed similar chromatin binding patterns (Fig. 1*D*). Binding profile of BALL closely mirrored that of CBP and H3K27ac at a majority of genomic regions. Importantly, most of these shared loci were also expressed in S2 cells, as highlighted by analysis of RNA-seq data (9) (Fig. 1*E*). About 3114 of 4577 (68%) genes, where BALL and CBP co-bound, were indeed found to be actively transcribed in S2 cells (Fig. 1*F* and Dataset S1). Comparison between top scoring DNA motifs to which BALL and CBP preferentially bind, revealed a distinctive similarity (Fig. 1*G*). Collectively, our data highlights that BALL substantially mimics the binding of CBP across the whole genome. These findings prompted us to investigate whether BALL and CBP biochemically interact with each other. To this end, co-immunoprecipitation (co-IP) experiments were performed using S2 cells that stably expressed FLAG-tagged BALL. Immunoprecipitation of BALL-FLAG resulted in precipitation of endogenous CBP as compared to empty vector control (Fig. 1*H*). To confirm this interaction, a reciprocal co-IP was performed using anti-CBP antibody, which resulted in immunoprecipitation of BALL-FLAG (Fig. 1*I*) as compared to mock IP. These results illustrate a hitherto unknown interaction between BALL and CBP, which may explain their co-occupancy at actively transcribed regions.

**Fig. 1.**
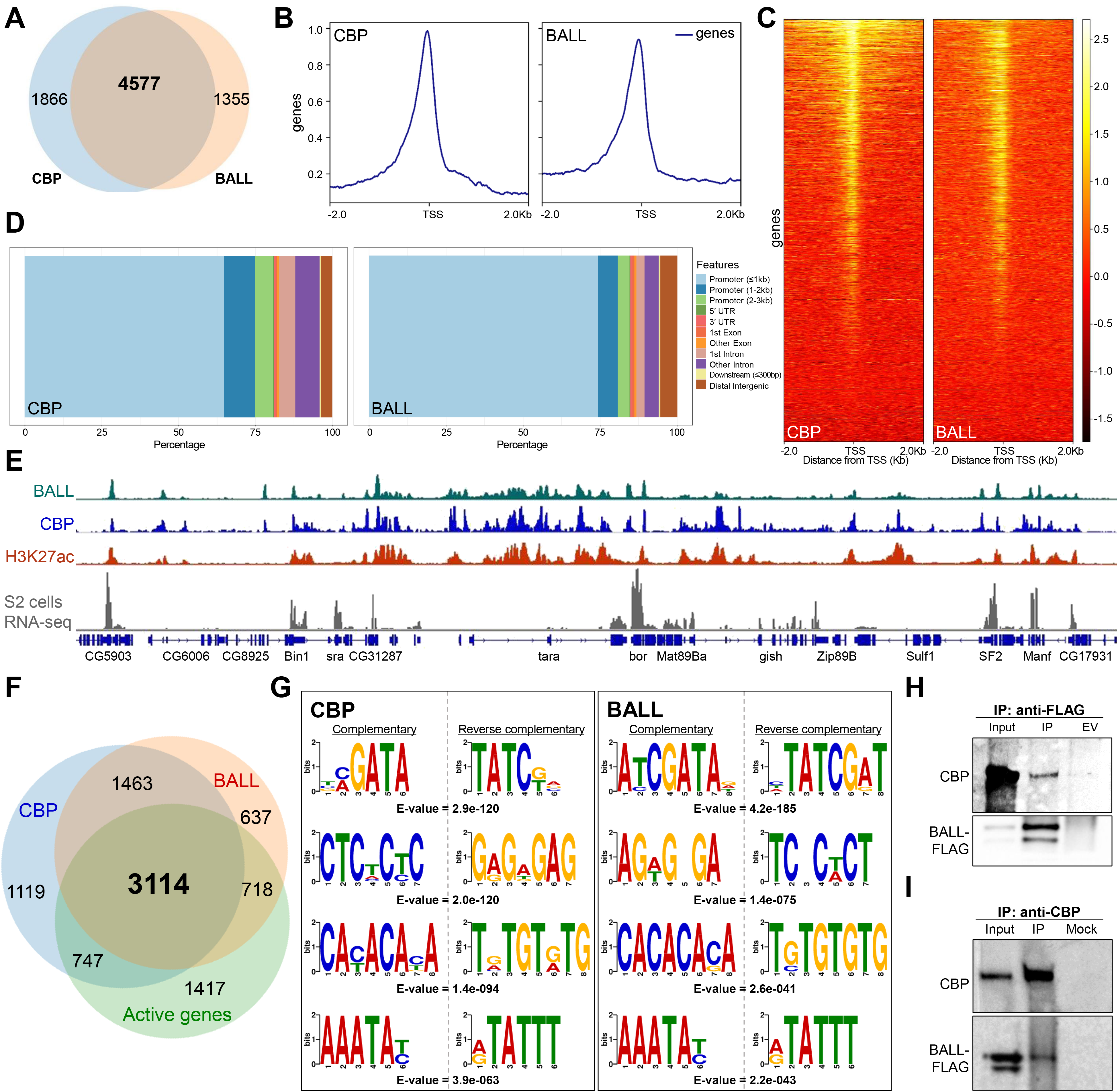
Genome-wide occupancy of BALL, CBP, and their biochemical interaction in *Drosophila* S2 cells. (A) Venn diagram depicting substantial overlap between CBP and BALL bound genes in S2 cells. Both BALL and CBP share 4577 binding sites at chromatin. (B, C) Comparison of genome-wide occupancy of CBP and BALL reveals an enormous similarity. Both CBP and BALL preferentially bind chromatin upstream to the TSS (B). Heat maps covering ±2 Kb region from TSS illustrate CBP and BALL occupancy across the genome (C). CBP and BALL mainly co-occupy the regions from TSS to 1Kb upstream. (D) Percentage distribution of CBP and BALL binding across different genome features shows a similar pattern where both CBP and BALL mainly reside in promoter regions. (E) Integrated genome browser view of the ChIP-seq data shows overlap between chromatin binding profiles of BALL (top panel), CBP (middle panel) and H3K27ac (lower panel). Genes co-occupied by BALL and CBP display active transcription as inferred by RNA-seq analysis of S2 cells (bottom panel). (F) Venn diagram depicting CBP and BALL enriched genes overlaid with actively transcribed genes in S2 cells. Majority of the genes that were co-bound by CBP and BALL were found to be transcriptionally active. (G) Four of the top five short DNA binding motifs of CBP, generated from MEME-ChIP database, show-striking similarity with the binding motifs of BALL. Each motif is presented with its respective reverse complementary motif and E-value. (H) Immunoblots of endogenous CBP and FLAG-tagged BALL after co-immunoprecipitation (IP) using anti-FLAG antibody in *Drosophila* S2 cells containing inducible BALL-FLAG transgene. As compared to IP from empty vector (EV) control cells, noticeable enrichment of CBP can be seen in IP using anti-FLAG antibody, which also resulted in enrichment of FLAG-BALL. (I) Immunoblots of endogenous CBP and FLAG-tagged BALL after co-immunoprecipitation (IP) using anti-CBP antibody in *Drosophila* S2 cells. BALL-FLAG is enriched after IP with anti-CBP antibody as compared to mock IP used as negative control. Enrichment of CBP validates successful IP.

Next, we investigated if depletion of BALL affects CBP mediated H3K27ac levels globally. Western blot analysis of cells where *ball* was knocked down, revealed 58% reduction in H3K27ac levels as compared to control cells treated with dsRNA against *LacZ* (Fig. 2*A*). To validate this finding *in vivo*, we generated homozygous *ball^2^* mitotic clones using the flp/FRT system. Immunostaining of imaginal discs with *ball^2^* mutant clones showed a drastic reduction in H3K27ac (Fig. 2B–E). Our results demonstrate that BALL positively contributes to CBP mediated H3K27ac. Since BALL is a histone kinase, it is imperative to investigate whether the impact that BALL exerts on H3K27ac is through the phosphorylation of neighboring H3 residues or due to the direct phosphorylation of CBP or its interacting proteins. Interestingly, VRK1 (Vaccinia related kinase 1), the mammalian homolog of BALL, was shown to phosphorylate CREB at serine 133 (10). This phosphorylation is known to enhance CREB interaction with CBP, leading to increased transcription of CREB target genes (11). The VRK1, CBP and CREB nexus may explain the regulation of CBP mediated gene activation but warrants further investigation.

**Fig. 2.**
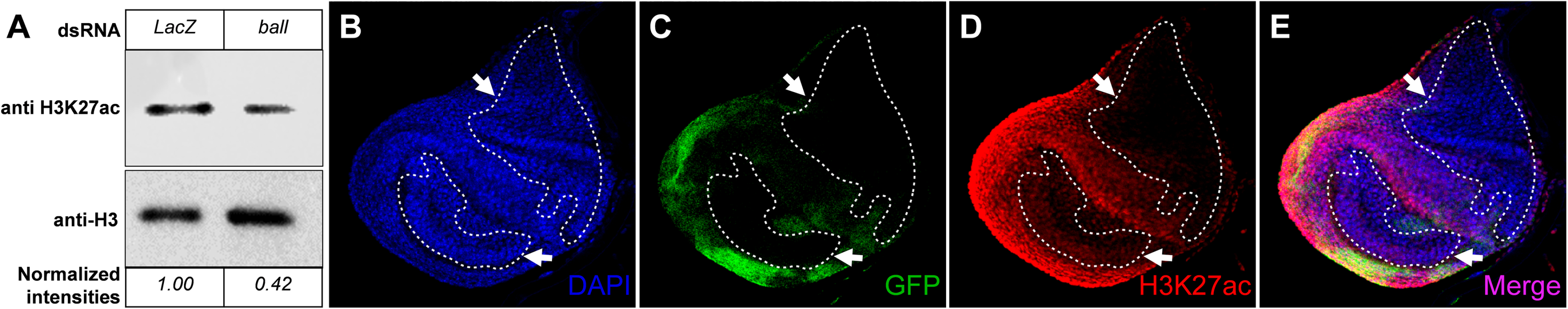
Depletion of BALL affects H3K27ac levels. (A) Western blot analysis of whole cells lysates from *Drosophila* S2 cells treated with dsRNA against *ball* exhibit reduced H3K27ac. Cells treated with dsRNA against *LacZ* served as control while total levels of histone H3 were used as a normalization control in immunoblot. Analysis of the relative intensities of H3K27ac signal, normalized against total H3 levels using ImageJ, showed 58% reduction in H3K27ac upon *ball* depletion. (B-E) Haltere imaginal discs containing mitotic clones of *ball^2^* were stained with GFP (C) and H3K27ac (D) antibodies. Mitotic clones, marked by the absence of GFP, showed drastic reduction of H3K27ac. Imaginal discs showed uniform DAPI staining (B). Mitotic clones are encircled and highlighted with arrows.

## Materials and Methods

Detailed materials and methods for ChIP-seq data analysis, molecular cloning, generation of BALL stable cell line, co-immunoprecipitation and mitotic clones for *ball^2^* are provided in the *SI Appendix*.

## Supporting information

Data Set 1

## Acknowledgements

We would like to thank Alexander M. Mazo for anti-CBP antibody and Alf Herzig for providing us *ball* flies for generating mitotic clones.

