## Supplementary Information for "Interplay between BALL and CBP maintains H3K27 acetylation on active genes in *Drosophila*"

**SI Appendix: Methods**

**ChIP-seq analysis**

ChIP-seq analysis was done as described previously (1). Briefly, the sequencing datasets were uploaded to the Galaxy web public server (2). Using Bowtie version 2, the ChIP-seq data was mapped to *Drosophila* genome (dm6) (3). Peak calling was performed using MACS version 2 with FDR (q-value) of 0.05 as cut-off and read extension of 200 bp (4). List of annotated peaks was extracted from peak file using ChIPseeker (5). Comparative heat maps of BALL and CBP were generated using deepTools, (bamComapre, computeMatrix and plotHeatmap) (6). MEME-ChIP - motif discovery, enrichment analysis and clustering on large nucleotide datasets (Galaxy Version 4.11.2+galaxy1) was utilized to generate DREME (Discriminative Regular Expression Motif Elicitation) output of BALL and CBP ChIP-seq data for de-novo motif discovery (2). ChIP-seq data of CBP, BALL and H3K27ac were taken from GSE64464 (7), GSE165685 (1) and GSE81795 (8), respectively. RNA-seq data of *Drosophila* S2 cells (GSM480160) (9) was mapped using TopHat Gapped-read mapper for RNA-seq data (Galaxy Version 2.1.1). Exon read count was calculated using htseq-count to generate list of active genes.

**Generation of BALL Stable cell line**

To generate stable cell line with inducible expression of FLAG tagged BALL, *w^1118^* embryos were used to prepare cDNA and amplify *ball* CDS. The *ball* CDS was first cloned in *pENTR-d-TOPO* entry vector (ThermoFisher Scientific) followed by sub-cloning in *pMTWHF* destination vector from DGVC (*Drosophila* Gateway Vector Collection) by setting up LR Clonase reaction following manufacturer’s protocol (ThermoFisher Scientific). The resulting *pMT-ball-FLAG* vector was transfected into S2 cells using Effectene transfection reagent (Qiagen) and a stable cell line was generated by following the manufacturer’s instructions (Qiagen).

**Co-immunoprecipitation**

For co-immunoprecipitation, S2 cells expressing FLAG-tagged BALL were harvested by centrifugation at 3000rpm for 5 minutes and washed twice with PBS. Cells were suspended in 1mL lysis buffer containing 140mM NaCl, 20mM Tris (7.4 pH), 1mM EDTA, 0.5% NP40, 10% glycerol, 0.2mM Na_2_VO_4_, pepstatin 0.5µg/mL, leupeptin 0.5µg/mL, aprotinin 0.5µg/mL and PMSF 1mM followed by incubation on ice for 30 minutes. The lysate was centrifuged at 14,000rpm for 15 minutes and supernatant was added to 40µL of M2-FLAG agarose beads (Sigma-Aldrich) which were pre-washed with lysis buffer. After one hour of incubation, beads were washed thrice with lysis buffer, resuspended in 30µL Laemili Loading buffer and heated at 95ᵒC for 5 minutes. IP samples were loaded on Novex precast Tris-acetate 3-8% gradient gels (ThermoFisher Scientific). For CBP IP, 1:1 Protein A and G Dyna beads (ThermoFisher Scientific) mixture was used and incubated overnight with anti-CBP antibody. Cell lysates were prepared, and incubated for one hour at 4ᵒC with the beads-antibody complex. After washing thrice with lysis buffer, samples were proceeded for western blotting as described above.

**Ex vivo knock down and mitotic clones**

Knock down of *ball* in D. Mel-2 cells was achieved as described previously (10). Briefly, D. Mel-2 cells were treated with 10 µg/mL of respective dsRNA for 4 days and total cell lysates were prepared in Laemili loading buffer before heating at 95ᵒC for 5 minutes. The mutant clones for *ball^2^* were generated using flp-mediated recombination (11). Fifty-five hours after egg laying, larvae with the genotype; *y^1^ w* P{ry^+^, hs-FLP}1/w*; P{neoFRT}82B P{Ubi-GFP}83/ P{neoFRT}82B e ball^2^* were given heat shock for one hour at 38ᵒC. Sixty-five hours after the heat shock, larval carcasses were inverted, fixed and labeled with antibodies against H3K27acand GFP. Immunostaining of imaginal discs was done as described previously (12). Images were acquired using Nikon C2 Confocal Microscope.

**Antibodies**

Antibodies used during this study are as following: rabbit anti-CBP (gift from Alexander M. Mazo, IP: 5µL, WB: 1:3000), mouse anti-GFP (Roche, 11814460001, IF: 1:50), mouse anti-FLAG M2 (Sigma Aldrich, WB: 1:2000), rabbit anti-H3K27ac (Abcam, Ab4729, IF: 1:75, WB: 1:2000), mouse anti-H3 (Abcam, ab10799, WB: 1:2000).
